# Bacterial, fungal and protist communities affect differentially in succession stage of karst soil

**DOI:** 10.64898/2025.12.15.694508

**Authors:** Li-Chun Mao, Gang Hu, Zhong-Hua Zhang, Cong Hu, Chao-Hao Xu, You-Huan Gan

**Affiliations:** School of Environmental and Life Sciences, Nanning Normal University, Nanning, China; Propaganda department, Nanning Normal University, Nanning, China

**Keywords:** Karst succession stage, Microbial communities, bacterial, fungal, protist

## Abstract

Karst soil microbial communities are major drivers of cycling of soil nutrients that sustain plant growth and productivity. Yet, a holistic understanding of the impact of different succession stage on the karst soil microbiome is still pooly understood. Here, we used a field experiment to investigate the loog-term consequences of changes in succession stage: rocky desertification(RD), grassland(GL), shrubs(SH), forest plantation(FP) and natural forest(NF) based on plant frequency on bacteria, protist and fungal communites. We showed that the shannon index of bacteria and fungi were higher in NF compared to GL, while protist shannon index was higher in SH that GL. The bacterial simpson_e index in NF and PF were higher than that in GL, SH and RD, howerer fungal and protist simpson_e index were similar across the five succession stages. Significant differences were observed in observed outs and phylogenetic diversity index in NF, SH were higher than GL, RD for fungi and protist, whereas bacterial observed outs and phylogenetic diversity index were relatively similar in five succession stage. The five succession stage differed markedly in the compositions of bacterial, fungi and protist communities. In bacterial, dominant phyla were Actinobacteriota, Proteoprotist, Acidobacteriota. Protist communities were dominated by Phragmoplastophyta, Arthropoda phyla. Regarding protist taxa, the dominant phyla were Ascomycota and Basidiomycota. Our findings provided evidence indicating that the microbial community compositions of RD, GL, SH, PF, and NF have their respective characteristics. These characteristics stem from differences in vegetation coverage, plant debris, root formation and other factors, leading to changes in the diversity and taxonomic group of bacteria, fungi and protists at different successional stages. Overall, this study provides a comprehensive perspective, showing the differences in microbial communities at different successional stages in karst soil.

## Introduction

Rocky desertification is one of the main causes of soil erosion in karst areas in Southwest China[1, 2]. The rocky desertification of land also threatens the multifunctionality of ecosystems and increases negative impacts on the environment, such as decreased carbon neutrality and nitrogen leaching losses[3-7]. Rocky desertification is usually related to changes in vegetation, which in turn can affect soil microbial communities[8-11]. As is well known, above ground vegetation affects underground microorganisms through factors such as the infiltration of organic carbon or changes in soil properties near plant roots[12-16]. In addition, Rocky desertification caused by grazing, logging can affect soil microbial communities[17, 18]. Due to the fact that soil microorganisms are the main participants in biogeochemical cycles, they are crucial for ecosystem functioning[19, 20]. Therefore, it is important to understand how microbial communities are affected and to what extent these reactions by succession stage of karst soil.

Soil microorganisms are vital components of the soil ecosystem, as their activities directly or indirectly be influenced by material cycling and energy flow, water-holding capacity, soil structure, and soil fertility[21-23]. Soil microorganisms such as bacterial, fungi, and protists play a critical role in realizing soil biodiversity, thereby influencing ecosystem functions[24-31]. For example, often overlooked protists are important in succession stage of karst soil as they control microbial populations through predation and influencing ecosystem functions such as nutrient cycling[32-34]. Especially, by preying on protist, phagotrophic protists are increasing the mobilization of bacterial nitrogen, which contributes to nitrogen mineralization[35]. This indicates the close interaction between soil microorganisms in different fields, which can affect soil function. Also, microorganisms are not just collections of independent populations, but form complex communities, direct and indirect interactions between taxonomic groups seem to play a crucial role in the assembly of microbial communities [36].Therefore, how soil microorganisms and their interaction mechanisms respond to succession processes in fragile ecological regions is crucial for maintaining soil nutrient cycling and preserving landscape integrity. However, research on this key focus within karst ecosystems is often overlooked.

Here, we investigated how the legacy effects of different stages of rocky desertification restoration affect the diversity, structure within and among bacterial, protist and fungal communities. For this purpose, we utilized the rocky desertification karst area in Apple County and established different succession gradients for rocky desertification restoration, including rocky desertification (RD), grassland (GL), shrubs (SH), forest plantation (FP) and natural forest (NF). We assume that at different stages of vegetation restoration in rocky desertification, changes in soil use will lead to different reactions among microbial communities, with the greatest differences between rocky desertification areas and nature forests; The communities of soil microbiota will gradually change along different succession stages of karst rocky desertification, with higher network complexity in forest systems, as nature forests support the coevolution of multi nutrient interactions within soil microbiota.

## Materials and Methods

### Experimental design and soil sampling

Soil samples were collected from a various successional stages in Pingguo, Guangxi province (107°15′ 42″E, 23°26′4″N), which is a typical karst area. The experiment was carried out in November 2024 and divided in five blocks. Each block comprises the following successional stages: rocky desertification (RD), glassland (GL), shrubs(SH), planted forest (FP) and natural forest(NF). Five blocks soil were collected randomly from each replicated plot using a soil corner of 20 cm depth and 5 cm diameter to obtain 6 samples from each treatment. The dominant vegetation of the RD was composed of *Arthraxon hispidus*, the GL was composed of *Heteropogon contortus, Cynodon dactylon*, the SH was composed of *Lantana camara, Vitex negundo, Alchornea trewioides*, the FP was composed of *Zenia insignis Chun*, the NF was composed of *Excentrodendron hsienmu*. Plant debris and roots were removed from soil samples, then which were homogenized and sieved at 4 mm.

### Assessment of microbial community composition and diversity

DNA was extracted from the 30 soil samples. The diversity and composition of bacterial, protist and fungal communities was analysed by 16S rRNA, 18S rRNA and ITS1 gene region using Illumina sequencing. The amplicons of the bacterial, protist and fungal were generated for DNA extracts. The V3-V4 hypervariable region of the bacterial 16S rRNA gene was amplified by polymerase chain reaction (PCR) using the primers 338F (5’-ACTCCTACGGGAGGCAGCA-3’) and 806R (5’-GGACTACHVGGGTWTCTAAT-3’)[37]. The V4 hypervariable region of 18S rRNA gene was amplified by PCR using the primers 547F (5’-GCAGTTAAAAAGCTCGTAGT-3’) and 967R (5’-TTTAAGTTTCAGCCTTGCG-3’)[38, 39]. Fungal ITS1 region was amplified using the primers ITS1F (5’-CTTGGTCATTTAGAGGAAGTAA-3’) and ITS2R (5’-GCTGCGTTCTTCATCGATGC-3’) primers[40]. The PCR products were detected from 1% agarose gels, then were purified using Agencourt AMPure XP Kit according to the manufacturer instructions.

### Sequencing and bioinformatic analysis

Sequence data from the 30 soil samples was analyzed after the sequencing raw datas were downloaded. First, utilize fastp for quality control on Fastq data. Adopt a sliding window strategy with a window size of 4, an average quality value of 20 and a minimum retained sequence length of 200, with fastq files automatically removing primer sequences. Second, quality checks were conducted. Further, concatenate reads, remove duplicates, filter out chimeras and generate representative sequences from post-quality control datasets. Using vsearch in qiime2 to de novo cluster the sequence after chimera filtering with a clustering threshold of 97% for 16S rRNA, 18S rRNA gene and ITS. Taxonomy was assigned using SILVA reference database for 16S and 18S and UNITE reference database for ITS. In this study, the number of clean reads ranged from 207750 to 289172 for bacteria, with an average of 251791; the number of clean reads for protist ranged from 185946 to 420498, with 240322 for average number; for fungal, the number of clean reads ranged from 211120 to 297822, with 264898 for average number. To avoid analytical bias caused by differences in sample data, random read-count smoothing is applied to each sample once sufficient sequencing depth is achieved. Typically, the lowest read count among the sequencing samples is selected as the baseline and the read counts for all samples are uniformly smoothed to this value. Bacteria, protist and fungal α-diversity index be analyzed using qiime[1],which data is typically trimmed to the minimum sample size in the dataset for computation during analysis. Using qiime calculated the β-diversity distance matrix to detect variations in the structure of microbial communities.

### Statistical analysis

The present study examined the effect of five succession stage on the community composition, α-diversity and β-diversity index for bacteria, fungi and protists. The differences in the microbial α-diversity index of bacteria, fungi and protists were tested using two-way t-test with a significance level *p*-value <0.05. Beta Diversity is a comparative analysis of microbial community composition across different samples, specifically examining the diversity of microbial communities between samples. We employed Principal Coordinates Analysis (PCoA) to visualize the differences among bacterial, fungal and protists communities, utilizing the Bray-Curtis distance etric. PERMANOVA is a nonparametric multivariate analysis of variance method based on the Bray-Curtis dissimilarity matrix and 9999 permutations are performed on the sample groupings. In the test results, R^2^ indicates the degree to which groupings explain the variation among samples and Pr < 0.05 indicates that the analysis is statistically significant. The computational process is implemented using the vegan package in the R language. By calculating the correlation index (SparCC) for features with an average relative abundance > 0.001%, we screened data with correlations > 0.6 and *p*-values< 0.05 to construct a correlation network. Different modules within the network were identified using the community network algorithm’s greedy algorithm.

## Results

### In succession stage of karst soil impacts soil microbial communites

To characterize the soil bacterial, fungal and protist communities under RD, GL, SH, PF and NF, we used high-throughput sequencing targeting the bacterial, protist and fungal communities. The shannon index of bacteria was highter in NF compared to GL. Protist shannon index was higher in SH compared to GL. Significant differences in shannon index were observed for the fungal, with NF being higher than GL and RD. The bacterial simpson_e index was higher in NF and PF compared to GL, SH, RD. Fungal and Protist simpson_e index were observed similarly across the five succession stages. The observed outs and phylogenetic diversity index of bacterial were similar across the five succussion stages. The observed outs and phylogenetic diversity index of protist and fungal were different across the five succussion stages. Protist observed outs and phylogenetic diversity index were higher in SH, PF, NF than GL, RD. Fungal observed outs and phylogenetic diversity index NF, SH were higher than GL, RD(Fig.1).

**Fig. 1.**
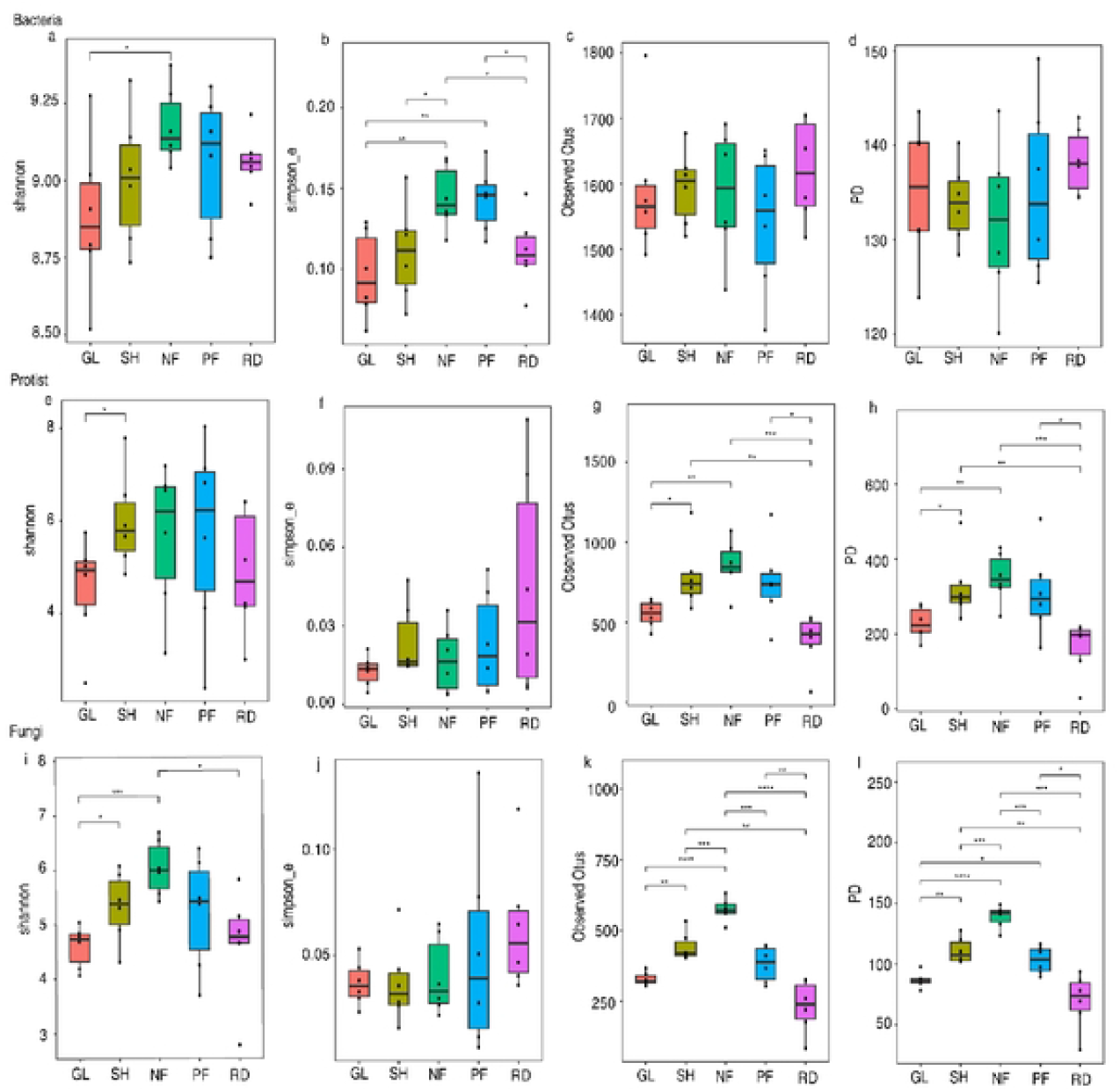
α-diversity of the microbial communities in RD, GL, SH, PF and NF. (a-d)bacteria shannon index, simpson_e index, observed species index, PD index; (e-h)protist shannon index, protist simpson_e index, observed species index, protist PD; (i-l)fungi shannon index, simpson_e index, observed species index, PD index. Significant differences is indicated by asterisks (\**p*<0.05,\*\**p*<0.01,\*\*\**p*<0.001).

Comparison of β-diversity using Principal Coordinates Analysis(PCoA) of Bray-Curtis distances showed a clustering of samples according to management with 51%, 27% and 30% of the variance explained by the first two axes of the PCoA for bacteria, protist and fungal communities respectively (Fig.2). Perm ANOVA confirmed significant differences in the structure of the microbial communities between seccession stage of karst soil(Fig.S1).

**Fig. 2.**
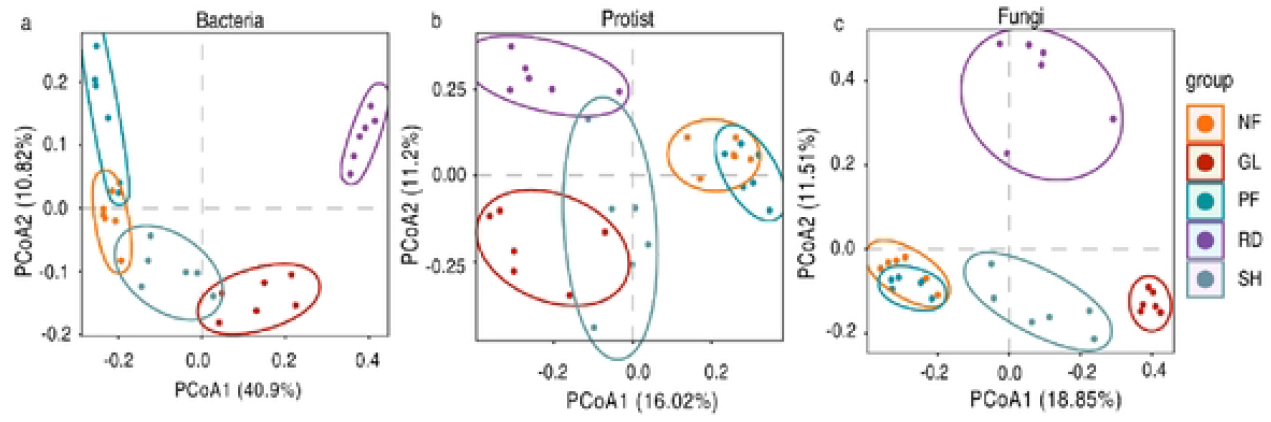
(a)PCoA of the Bray–Curtis distance matrices of bacterial 16S rRNA amplicons; (b)PCoA of the Bray–Curtis distance matrices of protist 18S rRNA amplicons; (c) PCoA of the Bray–Curtis distance matrices of fungal ITS1 amplicons.

### Identifying differentially abundant OTUs

Across all samples, the dominant bacterial phyla were p_Actinobacteriota(26%), p_Proteoprotist(19%), p_Acidobacteriota(20%). Protists were dominated by p_ Phragmoplastophyta(38%), p_Arthropoda(33%)and p_ Eukaryota_unclassified (17%), whereas p_ Ascomycota(90%)and p_Basidiomycota(35%)represented the most abundant fungal taxa (Fig.S2). The Venn diagram (Fig.3)shows the different variations and common characteristics of five succession stages. There are 975 bacteria OTUs, 194 protist OTUs, 68 fungal OTUs are shared in different succession stages. Bacterial OTUs of RD, PF, GL, NF and SH have 599, 401, 231, 199, 171 unique OTUs, respectively. Protist OTUs of RD, PF, GL, NF, SH have 721, 1240, 701, 1012, 963 unique OTUs, respectively. Fungal OTUs of RD, PF, GL, NF, SH have 286, 402, 196, 618, 325 unique OTUs, respectively. The ‘Upset’ plot shows the number of features that are common or different among samples in different succession stages.

**Fig. 3.**
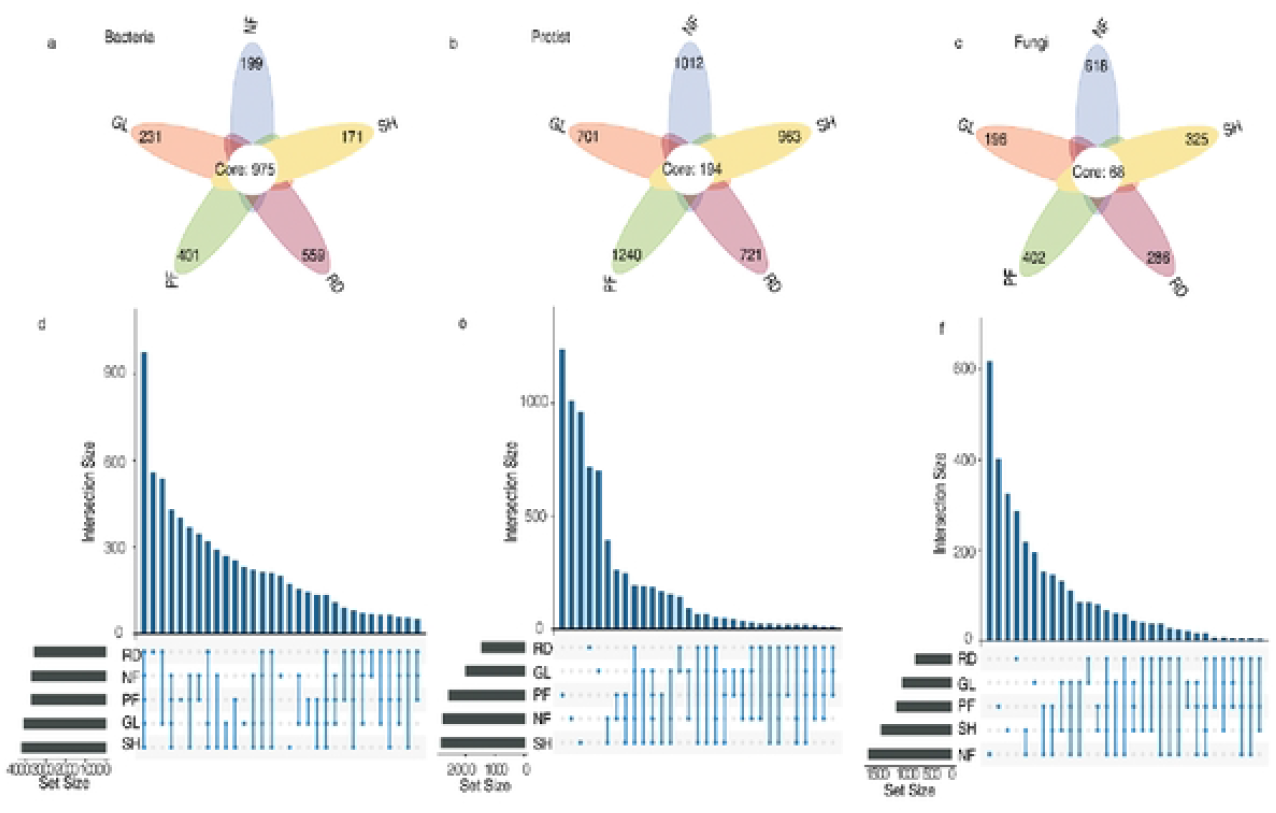
(a-c)The Venn diagram of bacterial, protist, fungi communities.(d-f)Upset plots showing the abundant OTUs for bacterial, protist, fungi.

### Co-occurrence networks within microbial groups are influenced by succession stages

To better understand how karst soil influences associations between bacteria, protist and fungi, we inferred co-occurrence networks according SparCC(weight>0.6, p<0.05) (Fig.4). For bacteria, Acidobacteriota, Actinobacteriota, Desulfobacterota, Entotheonellaeota, Gemmatimonadota, Methylomirabilota, Proteobacteria phylum are significantly correlated. Eukaryota_unclassified, Apicomplexa, Arthropoda, Ascomycota, Basidiomycota, Cercozoa, Chlorophyta, Chytridiomycota, Ciliophora, Cryptomycota, Nematozoe, Phragmoplastophyta, Rotifera phylum are significantly correlated for protist. For fungi, there are significantly correlated among Fungi_unclassified, Ascomycota, Basidiomycota phylum.

**Fig. 4.**
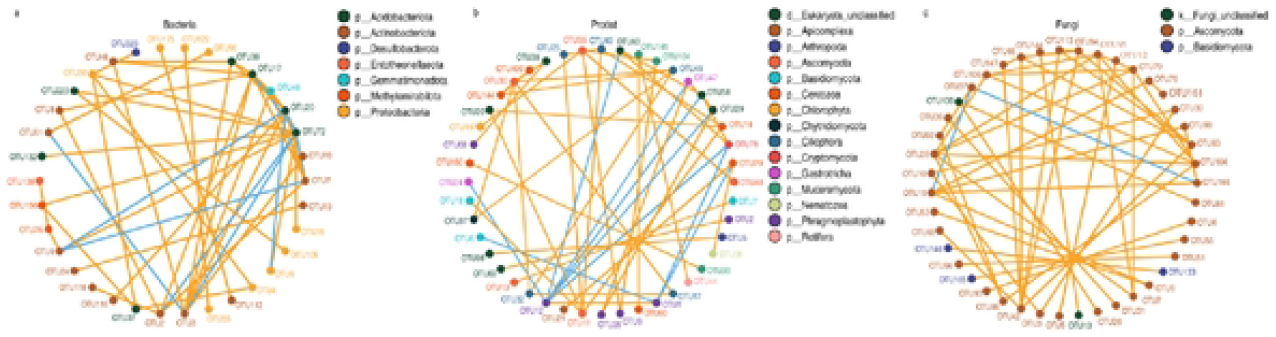
(a) Bacterial co-occurrence networks; (b) Protist co-occurrence networks; (c)Fungi co-occurrence networks. Each node represents a feature, with node size indicating the feature’s average abundance. Different colors denote the feature’s respective phylum. Edges represent correlations between features(orange indicating positive correlations and blue indicating negative correlations).

## Discussion

Investigations on the relationships between soil and microorganisms in karst ecosystems often focused on bacterial or fungal community. However, the research on protists how to influence the composition of the soil microbiome is scarce[41, 42]. Hence, it is highly imperative to adopt a more comprehensive approach to insight into the soil microbiome. Here, we compared and integrated analyses of soil bacterial, fungi and protist communities across different stages of karst vegetation succession.

### The diversity of bacterial, fungal and protist be effected by five succession stages

Our findings demonstrate that bacteria α-diversity showed differently. NF exhibited the highest Simpson_e values, with both NF and PF showing higher value than GL and RD. In line with previous studies, trees in natural and planted forests provided greater resources and energy for bacteria compared to desertified lands and grasslands, significantly enhancing bacterial diversity[15, 21, 43]; the extensive plant root systems in natural and planted forests partially mitigate soil erosion and nutrient leaching, thereby safeguarding soil quality[2, 43, 44]. These factors contribute to the higher bacterial diversity observed in natural and planted forests and explain to the rotation of plant communities in these systems that likely led to a greater diversity, ultimately increasing bacterial.

In terms of fungal diversity and richness, this study found that the Shannon index of NF was higher than that of GL and RD areas. This aligns with previous research indicating higher levels of plant detritus production increase the supply of organic substrates thereby increasing fungi diversity [45-47]. Alternatively, there are contrasting opinion that greater plant diversity increases the range of organic substrates entering soil, thereby increasing the number of niches according a greater array of heterotrophic fungi[48]. For instance, plant diversity enhanced soil fungal diversity and microbial resistance under plant invasion conditions; vegetation degradation have a key role in maintaining ecosystem functions and health throught obviously alters soil microbial diversities and hierarchical interactions [49, 50]. Furthermore, studies indicate that soil nutrient supply (e.g., plant root biomass) positively influences fungal diversity[51]. Thus, higher soil nutrient availability in NF and PF may contribute to their increased fungal diversity. NF exhibited greater fungal richness than GL, SH, PF and RD, attributing to fungi play a crucial role in decomposing plant residues and have a positive correlation between plant residues production and fungal diversity[48].

In terms of protozoan diversity and abundance, NF, PF and SH in this study exhibited higher diversity and abundance than GL and RD areas. This may be attributed to the larger plant root biomass in NF, PF and SH, which enables them to harbor greater microbial biomass, thereby increasing protozoan abundance[52, 53]. Soil protozoan diversity is influenced by soil pore water in heterogeneous environments[54]. Vegetation in NF, PF and SH enhances root biomass, stabilizes water availability and fosters stronger predation relationships between fungi and protozoa. Combined with increased soil carbon source availability and enhanced water stability, collectively explain the higher protozoan abundance in NF, PF and SH compared to grasslands and rocky desertification sites. We conclude that fungal abundance shows a significant correlation with protozoan presence[50], while bacterial abundance and diversity remain unrelated. However, some studies have reached opposite conclusions. For instance, research suggests that trends in prokaryotic diversity should highly correlate with bacterial diversity, with both reaching peak levels in eutrophic environments[55]. Some Complex symbiotic relationships exist between prokaryotic groups[56].

### The community composition be affected by five different succession stages

We showed that the β-diversity of all five domains is influenced by different successional soils, exhibiting significantly distinct community structures across the RD, GL, PF, NF and SH. This may stem from enhanced biodiversity driven by varying plant cover which in turn promotes biodiversity growth[57]. Instead, the smallest differences in community structure for all domains were observed between NF and PF, which suggests lasting effects of the plant. Microbial communities are important drivers of plant litter decomposition[58]. During vegetation succession, soil microbial communities exert a positive influence on soil microbial composition due to the impact of the nutrient environment[50]. Bacterial, fungal and protozoan communities in RD, GL, PF, NF and SH varied according to their respective succession stages. For instance, compared to other groups, NF, PF exhibited higher abundance of the proteobacteria phylum at the phylum level, This indicates that the survival of these categories relies more heavily on natural forest conditions, where organic matter sources enhance the presence of such microorganisms. Moreover, plant species can also affect differently soil microbial communities through a myriad of processes and create legacies that are detectable under subsequent plant communities, therefore also explaining the similarities between NF and PF. This study identified Actinobacteriota, Acidobacteriota, Proteobacteria as the dominant bacterial phylum across five successional stages, consistent with prior research indicating Actinobacteriota and Acidobacteriota dominate soil bacterial communities due to their competitive advantage in carbon utilization[59]. In terms of fungal microbial communities, the Ascomycota and Basidiomycota phyla occupy dominant positions across all five land use types. This finding aligns with observations in Guizhou’s karst regions[60], where the high calcium content of karst environments enhances the survival of both Ascomycota and Basidiomycota. Among protists, Phragmoplastophyta, Arthropoda, Eukaryota_unclassified dominated across the five successional stage. Their resilience to key soil parameters enables survival across broader soil types[61, 62]. Strong adaptation to moist environments resulted in significantly higher relative abundance of Phragmoplastophyta species in natural forest plots compared to other plots[63]. Here, enhanced soil structure facilitated root biomass growth, thereby stabilizing soil moisture content[57, 64, 65]. Arthropoda play a pivotal roles in nutrient cycling, decomposition, soil structure formation and soil biodiversity. The relative abundance of Arthropoda is higher in NF compared to other successional stage soil, indicating that their role as indivators of soil health due to their sensitivity to environment changes and enhance the activities of soil microorganisms[66, 67]. This finding is particularly significant given that such alterations in soil communities may subsequently influence plant communities[59, 65]. Due to inherent differences in ecological evolutionary dynamics, it is plausible that plant-soil feedback networks in NF may exhibit greater complexity and connectivity[68, 69]. As an ecological succession process, it can be inferred that plant-soil feedback networks in natural forests may form more intricate and interconnected structures[70-72].

Finally, we constructed a network model encompassing all five groups to obtain a comprehensive overview of different types of soil microbial communities. Given recent studies suggesting that ecosystem function may correlate with microbial network complexity, it is essential to examine how microorganisms across the five successional stages influence connectivity within soil microbial communities[73].

## Conclusions

By conducting a holistic microbiome investigation of bacteria, fungal and protist communities in different successional stages of karst landscapes, we showed that α-diversity of bacteria populations were affected across different succession stages. In complex forest systems, their diversity actually increased. However, we identified a clear difference in the structure and the composition of all communities in response to karst soil, in particular between NF and RD. The NF system led to more complex bacteria as well as inter-domain networks, which can have implication for the contribution of microbes to ecosystem multifunctionality. Inter-domain networks also revealed the predominant role of the protist as key taxa in soil microbiome networks across all karst soil. Future work need to validate the importance of protists in shaping soil microbial communities, directly through biotic interactions and/or indirectly through changes in abiotic factors.

